# Limits to the rate of information transmission through MAPK pathway

**DOI:** 10.1101/402750

**Authors:** Frederic Grabowski, Paweł Czyż, Marek Kochańczyk, Tomasz Lipniacki

## Abstract

Two important signaling pathways of NF-κB and ERK transmit merely one bit of information about the level of extracellular stimulation. It is thus unclear how such systems can coordinate complex cell responses to external cues. Here, we analyze information transmission in the MAPK/ERK pathway that features relaxation oscillations and responds to EGF by pulses of activated ERK. Based on an experimentally verified computational model of the MAPK/ERK pathway, we demonstrate that when input sequences of EGF pulses are transcoded to output sequences of ERK activity pulses, transmitted information increases nearly linearly with time. Moreover, the information channel capacity *C* (defined as the upper limit of information that can be transmitted over a sufficiently long time *t,* divided by *t*), is not limited by the bandwidth *B* = 1/τ, where τ ≈ 1 hour is the relaxation time. Specifically, when input is provided in the form of sequences of short binary EGF pulses separated by varying intervals that are multiples of τ/*n* (but are not shorter than τ), then for *n* = 2, *C* ≈ 1.39 bit/hour; and for *n* = 4, *C* ≈ 1.86 bit/hour. We hypothesize that the primary mode of operation of the MAPK pathway is to translate extracellular growth factor “bursts” into precisely timed intracellular ERK “spikes” of a predefined amplitude. Such pulse-interval transcoding allows to relay more information than the amplitude–amplitude transcoding considered previously for the ERK and NF-κB pathways.

**Author summary:** To coordinate their actions, cells communicate with each other by sending, receiving, and interpreting cytokine signals. Cells recognize chemical identity of signaling molecules and process quantitative and temporal properties of stimulation: amplitude, duration, or frequency. Previous studies indicated that the MAPK pathway transmits about one bit of information about the amplitude (concentration) of a stimulating cytokine, EGF. Sending more information may be enabled by temporal signal modulation. Here, we use the conceptual framework of information theory to support a hypothesis that the MAPK pathway, analyzed as a noisy information channel, reaches maximum information transmission rate when receiving sequences of EGF pulses that are transcoded to sequences of activity pulses of an effector kinase of the pathway, ERK. Since the pathway resetting time is about 1 hour, one could expect that—when sending EGF pulses of “0” or “1” amplitude every hour—the pathway is able transmit up to 1 bit per hour. We show, however, that when EGF pulses are separated by intervals that are not shorter than 1 hour and are multiples of 15 min (60, 75, 90 min, etc.), information rate can be nearly 2 bits per hour. We hypothesize that high information rate is necessary to control cell proliferation and motility.

## Introduction

Cells recognize and respond to chemical signals in order to adapt to changing external conditions and coordinate their actions at a supercellular level. Not only chemical identity of an incoming signal but also its quantifiable features such as amplitude or duration are usually relevant for eliciting a proportionate physiological cell response. The amplitude of an input signal can be translated into the amplitude or duration of activity of an effector molecule such as a transcription factor or a small-molecule secondary messenger. It has been shown in the NF-κB system that the response in a population of cells is proportional to the product of both signal amplitude and its duration [1]. Some signaling pathways that exhibit oscillatory behavior at the single-cell level are able to transcode input amplitude to the frequency of effector pulses [2]. Such capacity has been recently demonstrated in the mammalian mitogen-activated protein kinase (MAPK) pathway, where in response to extracellular EGF (input) of constant amplitude, ERK (effector) is activated in pulses with constant amplitude but frequency and pulse duration determined by concentrations of EGF [3]. In the MAPK system, the amplitude-to-frequency transcoding (also termed modulation in engineering sciences) is enabled by a particular topology of feedback loops and associated timescales [4].

Molecular interpretation of pulsatile (or more complex) temporal codes of effector molecules is an area of active research [5–7]; however, less attention has been devoted to the characterization of responses of signaling systems challenged with time-varying inputs. In controlled conditions of microfluidic systems it has been demonstrated that both the NF-κB and the MAPK/ERK pathways respond in a pulsatory manner to periodic pulses of cytokines, TNF and EGF, respectively [8,9]. Experimental and computational analysis of responses to pulsating inputs has shown that the ability of the NF-κB system to respond with a high fidelity to a series of TNF pulses is inherently limited by the relaxation time τ associated with an oscillations-generating negative feedback [10,11]. Information transmission through both the NF-κB and the MAPK/ERK pathways has been analyzed within the generic framework of information theory [12,13], revealing that these pathways are able to transmit about 1 bit of information about a constant stimulus concentration [14–17]. This estimate increases slightly when the trajectory of the effector is sampled at several time points [16].

In this work, we present a computational analysis of information relay within an experimentally verified model of the MAPK/ERK pathway [4], where a fast positive feedback loop (involving RAS and SOS) is nested within a slow negative feedback loop (involving ERK and SOS; see Fig 1A). Such network topology allows both to transcode constant EGF stimulation to periodic ERK activity pulses and to transcode analogous EGF pulses to nearly digital pulses of ERK activity. Based on the considered model, we approach the following representation problem: What is the signal transcoding strategy that allows to transmit maximum amount of information through a given transmission channel in a given time span or to achieve the highest information transmission rate in the long term? We propose to code input information in the form of sequences of short digital EGF pulses. Then, we estimate from below the maximum transmission rate with respect to the noise strength. In particular, we demonstrate that when input information is encoded in intervals between subsequent EGF pulses, information channel capacity, computed as the maximum mutual information rate, may exceed channel bandwidth, *B* = 1/τ. By theoretical and numerical analysis of transmission of binary EGF sequences with varying inter-pulse intervals (not shorter than the relaxation time τ of about 1 hour), we demonstrate that the MAPK/ERK pathway is able to transmit more than 1 bit per hour.

**Fig 1.**
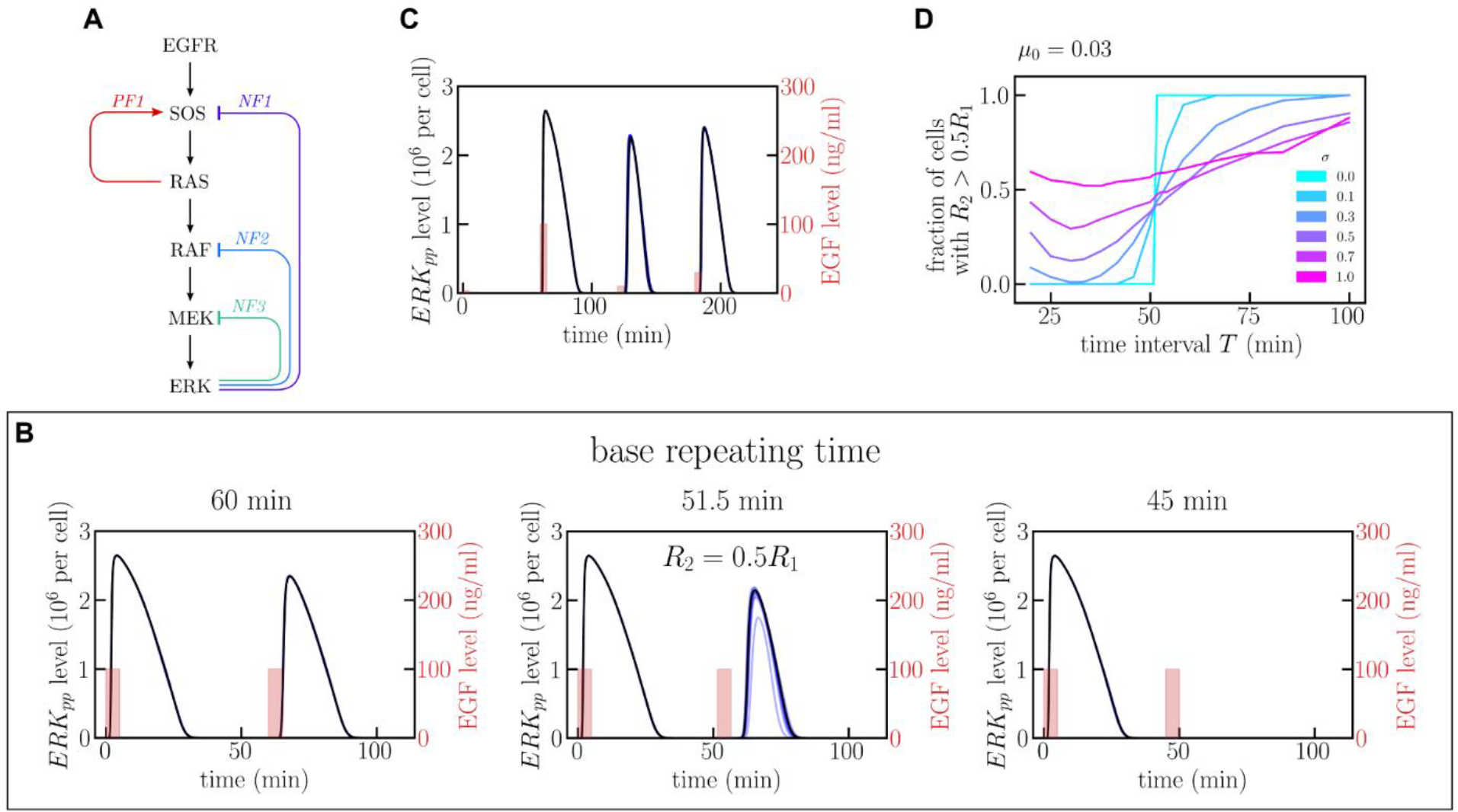
Diagram of the MAPK pathway and analysis of ERK responses to repeated EGF pulses. (A) Schematic diagram of the MAPK pathway. As analyzed in [4], PF1 and NF1 lead to relaxation oscillations, whereas NF2 and NF3 shape time-profile of the pulse of active ERK (*ERK*_*pp*_). (B) *ERK*_*pp*_(*t*) profile in response to two 5 min-long 100 ng/ml EGF pulses separated by *T* = 45, 51.5, 60 min. Black line: deterministic model trajectory; blue lines: 10 stochastic trajectories (visible only for *T* = 51.5 min). (C) *ERK*_*pp*_(*t*) profile in response to four EGF pulses repeated every 60 min with successive amplitudes 3, 100, 10, 30 ng/ml. (D) Fraction of simulated cells for which *R*_2_ > 0.5*R*_1_, as function of the time interval *T* between EGF pulses. The fraction is calculated based on at least 500 independent numerical simulations (number of simulations for *T* close to τ was up to 5000) performed assuming a lognormal distribution LogN(µ_*i*_, σ^2^) of protein levels (and first order reaction coefficients) with mean µ_*i*_ equal to the default value of a given variable, and six different values of σ.

## Results

### ERK responses to the repeated EGF pulses of different frequencies and amplitudes

The ability of a cell to respond to EGF pulses depends on their amplitude, duration, and pulsing frequency. We focus on relatively short pulses of 5 min. In Fig 1B we show trajectories resulting from one deterministic (black line) and ten stochastic (blue lines) simulations of the MAPK model in response to two EGF pulses, both of amplitude 100 ng/ml, separated by three different time intervals *T* of 45, 51.5, and 60 min. Let *R*_*i*_(*T*) denote the integrated ERK response after *i*th EGF pulse:

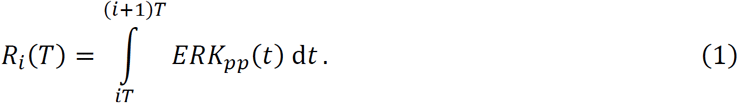

Since the ERK_pp_ pulse lasts about 30 min, for *T* > 30 min, *R*_1_(*T*) = const =: *R*_max_ with the numerically calculated *R*_max_ = 2.65×10^9^ molecule×s. Let us consider the integrated ERK responses to the first and the second EGF pulse. For *T* = 45 min, the response to the second pulse, *R*_2_, is negligible with respect to *R*_1_, *R*_2_(45)/*R*_1_(45) ≪ 1, while for the time interval *T* = 51.5 min, *R*_2_(51.5)/*R*_1_(51.5) = 0.5. For *T* = 60 min, *R*_2_ = 2.00×10^9^ molecule×s is comparable to *R*_1_. This suggests that there is a threshold τ = 51.5 min above (below) which the system can (cannot) be activated in response to the next pulse. It is of note that this threshold time (relaxation time) τ is very close to the minimal period of oscillations under constant EGF stimulation observed experimentally [3].

In Fig 1C, we analyze the ERK activity time-profile in response to EGF pulses repeated every 60 min, with successive amplitudes 3, 100, 10, 30 ng/ml. While there is no response to the 3 ng/ml pulse, the responses to the remaining three pulses have comparable amplitudes, even though the 10 ng/ml pulse follows the much stronger 100 ng/ml pulse, which has inhibitory effect. Overall, the examples presented in Fig 1B and Fig 1C show that the MAPK system exhibits nearly all-or-nothing ERK responses to the pulsatile simulation. We thus expect that in the case of pulsatile stimulation, information is not coded by the level of EGF, but by presence or absence of an EGF pulse that exceeds some threshold. Consequently, in the further analysis we restrict ourselves to 5 min-long EGF pulses of amplitude 100 ng/ml.

We notice that the stochastic trajectories in Fig 1B and Fig 1C are nearly indistinguishable from the deterministic one, except for trajectories with *T* = 51.5 min in Fig 1B. This is a consequence of a large number of reacting molecules at each step of the pathway, which makes the intrinsic noise of signal processing negligible. When comparing model predictions with experiment we found that to reproduce experimentally observed heterogeneity of single cell responses, one has to include extrinsic noise, i.e., assume that the levels of the MAPK pathway components vary among cells [4].

Since intrinsic noise was found negligible, henceforth we will use a deterministic approximation (ordinary differential equations), which substantially speeds up numerical simulations. In order to analyze how extrinsic noise influences the transmitted information, we will consider two noise sources:

A. Cell-specific noise associated with uncertain levels of pathway components (in individual simulated cells) and with pseudo-first order rates of dephosphorylation reactions (by implicit phosphatases). We will assume that these variables follow the lognormal distribution LogN(µ_*i*_, σ^2^) with median µ_*i*_ equal to the default value of an *i*th parameter.
B. Additive noise associated with ERK activation mediated by pathways that are not included in the model. For the additive noise we will also assume a lognormal distribution LogN(µ^*^, σ_0_^2^) with σ_0_= 1 and µ^*^ = µ_0_×*R*_1_×*T*/(60 min). For most of the analysis we take µ_0_ = 0.03, which is equivalent to the assumption that about 3% of ERK activity results from the additive noise.

The parameters σ and µ_0_ that characterize the strength of the cell-specific and the additive noise will be varied to show how they influence transmitted information. Details of numeric simulations are given in Methods.

The MAPK channel information capacity (or simply bitrate) is controlled by the maximal frequency of EGF pulses that can induce ERK activation. For Fig 1D we perform similar simulations as for Fig 1B, but accounting for the extrinsic noise. We compute the fraction of cells for which the response to the second pulse is significant with respect to the first pulse, i.e., *R*_2_ > 0.5*R*_1_, as a function of time interval *T* between EGF pulses, for several values of σ (keeping µ_0_ = 0.03). As expected, for σ = 0, all cells exhibit significant responses to the second pulse when *T* > τ and none of them exhibit such responses when *T* < τ. For small and moderate noise, σ = 0.1 and σ = 0.3, to assure significant responses to the second EGF pulse, *T* must be substantially larger than τ; additionally, even for *T* < τ, a fraction of cells exhibits significant responses to the second EGF pulse. For larger noise, σ ≥ 0.5, regardless of *T,* there are cells that exhibit significant response to the second pulse and cells that fail to respond. In summary, based on the above analysis, we may expect that for zero or small noise, EGF pulses separated by intervals *T* > τ will lead to a significant ERK activation, whereas for larger noise, EGF pulses will be missed in some cells even for *T* > τ, but some other cells will show responses for *T* < τ.

### ERK activation in response to sequences of three EGF pulses

In Fig 2 we show responses to eight sequences of EGF pulses. We consider three base repeating times *T*_*i*_ (time intervals between repeated pulses) of 60, 30, or 20 min. In the following convention the sequence ‘101’ for *T* = *T*_1_ = 60 min denotes a sequence of three EGF pulses of successive amplitudes: 1 (in dimensional units 1×100 ng/ml) at *t* = 0 min, 0 (a “pseudo-pulse”) at *t* = 60 min, and 1 at *t* = 120 min. For *T* = *T*_2_ = 30 min, the same sequence denotes pulses of amplitude 1 at *t* = 0 min and *t* = 60 min, and a pseudo-pulse at 30 min.

**Fig 2.**
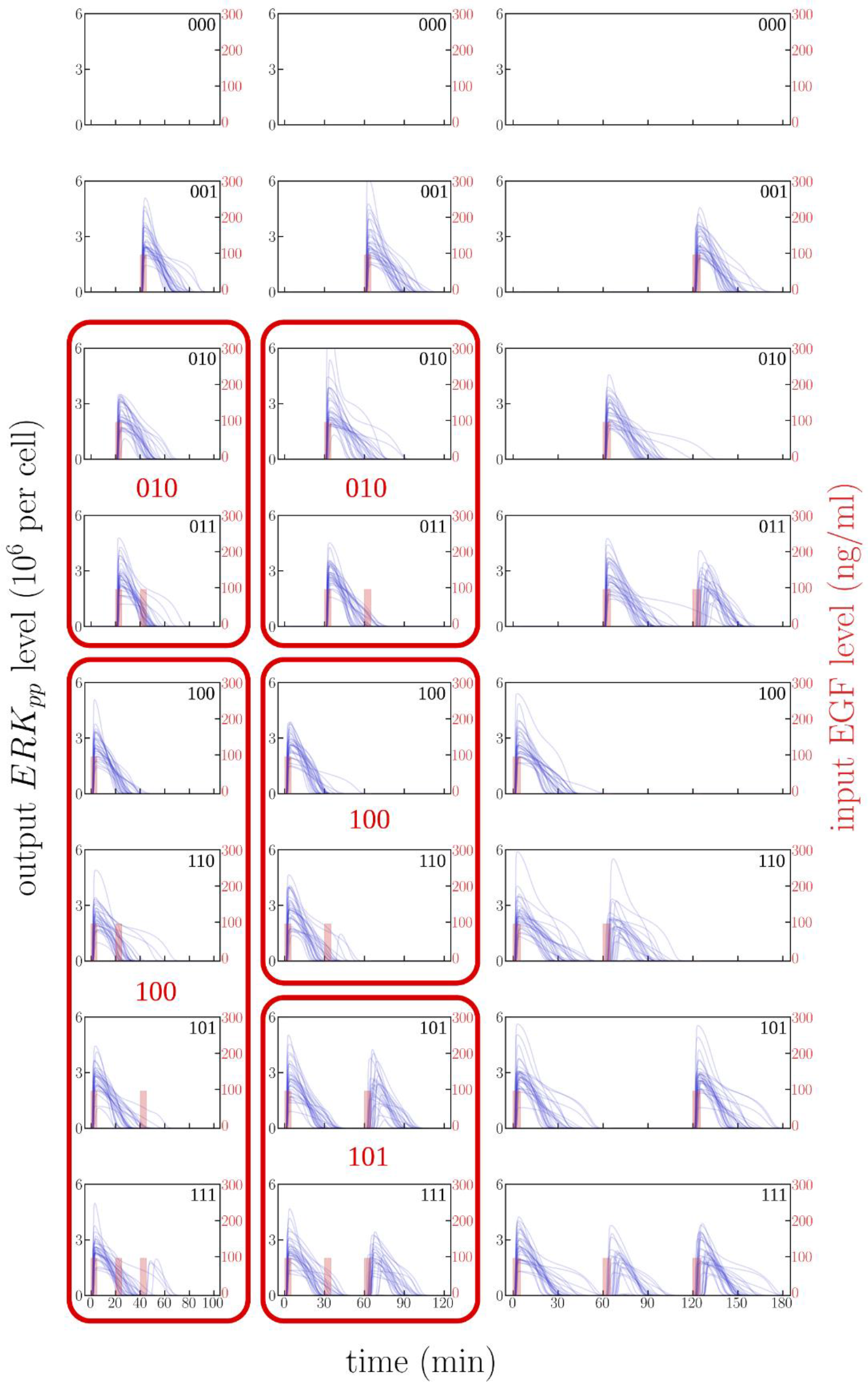
Responses to 8 possible series of 3 pulses having amplitude of either ‘0’ (“pseudo-pulse”), or ‘1’ (100 ng/ml in dimensional units). For each base repeating time *T*_*i*_ (60, 30, or 20 min) and each pulse sequence, 30 independent numerical simulations were performed with σ = 0.3, µ_0_ = 0.03. For *T* = 30 min and *T* = 20 min, the sequences transmitted to the same output sequence are grouped and encircled in red. First element of each group is accurately transmitted and will be referred to as a group representative.

For illustration purposes, times *T*_*i*_ are chosen so that *T*_1_ > τ, *T*_2_ < τ < 2*T*_2_, and 2*T*_3_ < τ < 3*T*_3_. For each of the eight sequences and each *T*_*i*_, we perform 30 simulations (each corresponding to a single cell) with σ = 0.3, µ_0_ = 0.03; the resulting trajectories of *ERK*_*pp*_(*t*) are shown in Fig 2. Since the noise is moderate, one can expect that for *T* = 60 min most cells will respond to all EGF pulses (by looking closely one can spot that only one and two cells show a very weak response to the second pulse in sequences ‘110’ and ‘111’, respectively). One can thus say that for *T* = 60 min almost all EGF sequences are transmitted properly.

For *T* = 30 min, cells are not able to properly transmit EGF sequences ‘011’, ‘110’, and ‘111’. Sequence ‘011’ leads only to a single ERK_pp_ pulse – in essence producing the same response as sequence ‘010’. We will say that these two input EGF sequences lead to a ‘010’ ERK_pp_ response. Similarly, EGF sequence ‘110’ is interpreted as ‘100’, whereas EGF sequence ‘111’ is interpreted as ‘101’. In the last case – because the second pulse does not elicit ERK activation – the third pulse can be transmitted. Overall, for *T* = 30 min, one can distinguish two EGF sequences, ‘000’ and ‘100’, that are accurately transmitted, and three groups {‘010’, ‘011’}, {‘100’, ‘110’}, and {‘101’, ‘111’} (grouped in red frames in Fig 2) that are transcoded respectively to ERK_pp_ sequences ‘010’, ‘100’, and ‘101’. The element of the group which is accurately transmitted (i.e., here ‘010’, ‘100’, and ‘101’) will be referred to as the group representative. For *T* = 20 min, the analysis is analogous: there are also two sequences that are accurately transmitted, but only two groups: one containing 2 EGF sequences, the other containing 4 EGF sequences. Because an ERK_pp_ pulse at the first position inhibits signal transmission for about 50 min, EGF sequences ‘100’, ‘110’, ‘101’, and ‘111’ are all transcoded to ‘100’.

Transmitted information is limited from above by log_2_(*K*), where *K* is the number of distinct output sequences. In our case, *K* = 8 for *T* = 60 min, *K*= 5 for *T* = 30 min, and *K* = 4 for *T* = 20 min. Thus, transmitted information (or mutual information, MI) is limited from above by respectively: log_2_(8) = 3, log_2_(5) = 2.32 and log_2_(4) = 2. The estimated bitrate, i.e., MI/(3*T*_*i*_), equals respectively 1 bit/hour, 1.55 bit/hour, and 2 bit/hour. As we can see, the bitrate is highest when the base repeating time is 20 min, which, of note, is only a fraction of the relaxation time τ. We will see that this observation remains true also for longer sequences and that such simple theoretical estimates agree well with numerical values obtained for small noise.

### Quantitative estimation of transmitted information

In order to estimate bitrate numerically we will calculate mutual information between input and output sequences. The inputs *x* ∈ *X* are EGF binary sequences of length *L*. We have performed simulations for *L* = 8 and *L* = 6. For each input sequence we performed *M* = 1000 simulations obtaining *M* response vectors *y* = (*R*_1_, …, *R*_L_) ∈ *Y*, where *R*_*i*_ are defined as in Eq. (1) with the additive noise included. These simulations probe a (continuous) output probability distribution *Y*. The analysis of short sequences (*L* <6 is based on truncated simulation data obtained for *L* = 6. In this way we gained larger statistics; for example, for *L* = 2 (Fig 3), we have 16,000 simulation for each of four binary input sequences.

**Fig 3.**
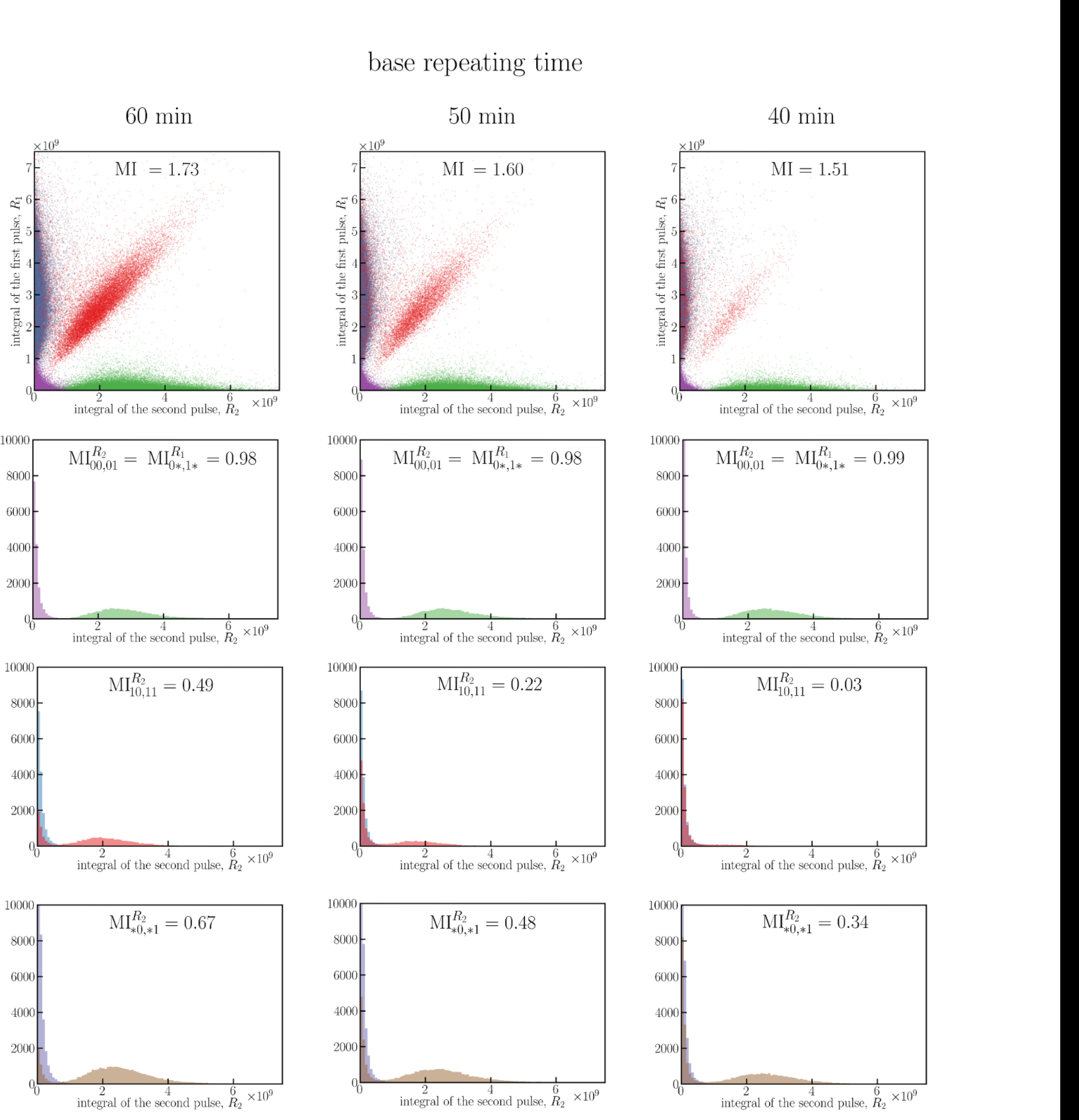
ERK responses to four binary sequences: 00, 01, 10, and 11 for three base repeating times. First row: scatter plots of *R*_1_, *R*_2_ for all sequences (purple 00, green 01, blue 10, red 11), based on 16,000 simulations for each sequence. Second row: histograms of *R*_2_ when the first pulse has amplitude 0 (purple 00, green 01). Third row: histograms of *R*_2_ when the first pulse has amplitude 1 (blue 10, red 11). Fourth row: histograms of *R*_2_ independently of the first pulse amplitude (blue *0, brown *1). In all scatter plots, MI denotes the mutual information calculated for two-dimensional distributions; in all histograms, MI denotes the mutual information calculated based on two shown one-dimensional distributions.

In our case the domain of input is discrete, so mutual information can be expressed as

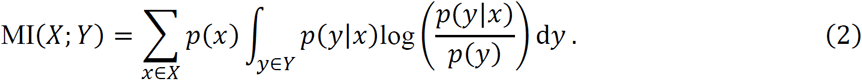

To calculate MI, one has to assume or determine input sequence probabilities *p*(*x*). We proceed as follows:

1. assume that all input sequences have equal probabilities, or
2. assume that all group representatives have equal probabilities, while the other sequences have zero probabilities, or
3. calculate input probabilities that maximize MI.

Description of the numerical approach used to estimate MI is provided in Methods.

### Numerical estimation of the transmitted information in sequences of two pulses

In Fig 3 we analyze the ability of the system to resolve binary sequences ‘00’, ‘01’, ‘10’, and ‘11’ for three base repeating times *T*_*i*_ = 60, 50, and 40 min, for moderate noise (σ = 0.3, µ_0_ = 0.03). In the first row of Fig 3 we present scatter plots in the (*R*_1_, *R*_2_)-plane showing responses to the four binary sequences with corresponding MI values. The scatter plot for *T* = 40 min shows that for this time interval the system is essentially able to transmit 3 input sequences, ‘00’, ‘01’, and ‘10), while the sequence ‘11’ produces almost the same output as sequence ‘10’. Consequently, MI = 1.51 < log_2_(3) ≈ 1.58. For *T* = 50 min and *T* = 60 min, the system distinguishes ‘10’ and ‘11’ sequences to a certain extent and MI values (equal respectively 1.60 and 1.73) exceed log_2_(3) but are still substantially lower than the upper bound Mi_max_ = log_2_(4) = 2.

We further investigate the system’s inability to transmit four sequences by detailed analysis of responses to the second pulse. When the amplitude of the first pulse is 0, then, regardless of *T*_*i*_, the system almost perfectly distinguishes between amplitudes 0 and 1 of the second pulse (see Fig 3, histograms in the second row). The value 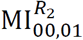 (given on each histogram in the second row) is calculated between the inputs *X* = {‘00’, ‘01’} (assuming that both sequences are equiprobable), and the output to the second pulse, *R*_2_. For each *T*_*i*_, 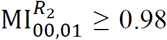 (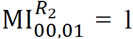 means that both inputs are perfectly distinguishable). In the case when the amplitude of the first pulse is 1, the ability to distinguish between amplitudes 0 and 1 of the second pulse is much lower (Fig 3, histograms in the third row): for *T* = 60 min, 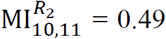 for *T* = 50 min, 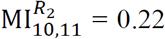 and for *T* = 40 min, 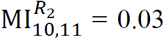 the nearly zero value for *T* = 40 min is a consequence of the fact that the system cannot respond to two EGF pulses with a time interval of 40 min. In the fourth row in Fig 3 we show histograms of the marginal probability distributions of responses to the second pulse. The 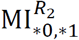 values shown in these histograms measure the ability to distinguish between amplitudes 0 or 1 of the second pulse when the first pulse is unknown.

Let us notice that two-dimensional MI is greater than the sum of unidimensional MIs, 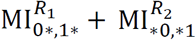 Here 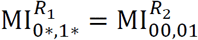, because the response to the second pulse in the case when the first pulse has amplitude 0 is identical to the response to the first pulse – regardless of the amplitude of the second pulse. This implies that information that can be inferred from two pulses together is higher than information inferred from the first and the second pulse separately (i.e., when the interpretation of the second pulse ignores information inferred from the first pulse). This indicates that a receiver with memory, that can record and analyze full response to an EGF sequence, can infer more information than a memoryless receiver that processes each pulse individually.

In the considered case, information about the response to the first EGF pulse allows to deduce more information from the second pulse. The response to the first pulse allows to almost unambiguously infer EGF input. Information about the second EGF pulse (knowing the response to the first pulse) is 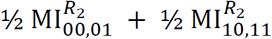, which exceeds 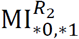. We notice that the overall MI is well approximated by 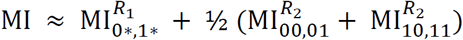 which implies that information transmitted in the sequence of two pulses is equal to information inferred from the first pulse plus information inferred from the second pulse knowing the amplitude of the first pulse. Unfortunately, this way of calculating the overall MI for longer sequences is impractical, and we will calculate MI based on multidimensional distributions of outputs to sequences of multiple pulses.

### Theoretical versus numerical estimation of information carried by pulse sequences

To estimate MI for small noise, one can express it in terms of entropy and conditional entropy:

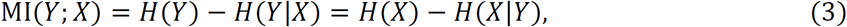

where *H*(*X*) and *H*(*Y*) are the entropies of input and output, *H*(*Y*|*X*) is the conditional entropy of output given input, while *H*(*X*|*Y*) is the conditional entropy of input given output. First, we note that this formula implies that MI is limited by the input entropy *H*(*Y*) as well as the output entropy *H*(*X*). In the simplest case of *T* > τ, all EGF sequences are properly transmitted (transcoded) to ERK sequences, and thus *H*(*Y*|*X*) = *H*(*X*|*Y*) = 0 (which means that the knowledge of input is sufficient to deduce output and, conversely, knowledge of output is sufficient to infer input). Thus, for *T* > τ we have MI(*Y*; *X*) = *H*(*Y*) = *H*(*X*) = *L* (where, recall, *L* is the length of binary input sequences). In this case, the bitrate is equal to the base repeating frequency Φ = 1/*T*. In the case of *T* < τ, MI(*Y*; *X*) = *H*(*X*) can be reached when one restricts input sequences to a subset that can be unambiguously transmitted, i.e., to group representatives (as defined for exemplary *L* = 3 in Fig 2).

Let us now consider the case of *L* = 4, *T* = 30 min. The input sequences form 8 groups: {‘0000’}, {‘0001’}, {‘0010’, ‘0011’}, {‘0100’, ‘0110’}, {‘0101’, ‘0111’}, {‘1000’, ‘1100’}, {‘1001’, ‘1101’}, and {‘1010’, ‘1110’, ‘1011’, ‘1111’}. In each group, input sequences are indistinguishable, so *K* = 8.The maximum mutual information is log_2_(8) = 3 and is achieved when probabilities of all group representatives (first sequence in each set) are equal to 1/8. The same maximal MI value is achieved when in each group the sum of all probabilities is equal to 1/8. In this case, one also learns 3 bits about input as knowing the output sequence allows one to infer from which of the 8 equiprobable groups the input signal originates. Unsurprisingly, by maximizing MI numerically in the case of small noise (σ = 0.1, µ_0_ = 0.03) we obtain almost equal probabilities of each group of input sequences (see Table 1, last column).

**Table 1.** Information transmitted in *L* = 4 pulses with intervals *T* = 60, 30, 20, 15 min. Comparison of numerical and theoretical estimates for small noise, σ = 0.1 and µ_0_ = 0.03.

| $T$<br>(min) | Theoretical MI value | | Numerical MI value, $\sigma = 0.1$ and $\mu_0 = 0.03$ | | | Maximum and minimum values of group probabilities together with the group representative.<sup>a</sup> |
| --- | --- | --- | --- | --- | --- | --- |
| | equal probabilities of input sequences | maximized $\log_2(K)$ | numerically maximized | equal probabilities of all group representatives | equal probabilities of all input sequences | |
| 60 | $\log_2(16)=4$ | $\log_2(16)=4$ | 3.93 | 3.93 | 3.93 | '0001': 0.065; '0110': 0.060 |
| 30 | $4-9/8=2.875$ | $\log_2(8)=3$ | 2.96 | 2.96 | 2.82 | '0010': 0.129; '1010': 0.120 |
| 20 | $4-13/8=2.375$ | $\log_2(6)=2.58$ | 2.55 | 2.55 | 2.33 | '0100': 0.171; '1001': 0.161 |
| 15 | $4-17/8=1.875$ | $\log_2(5)=2.32$ | 2.32 | 2.32 | 1.89 | '1000': 0.202; '0000': 0.195 |

<sup>a</sup> Data obtained via numerical MI maximization.

Let us notice that by assuming that all 16 input sequences have equal probabilities, we obtain a somewhat lower value of MI(*Y*; *X*) = *H*(*X*) – *H*(*X*|*Y*). Now, *H*(*X*) = log_2_(16) = 4, while *H*(*X*|*Y*) = (2/16)log_2_(1) + (5/8)log_2_(2) + (1/4)log_2_(4) = 9/8, which gives MI(*Y*, *X*) = 23/8 < 3. The calculations for *T* = 20 min and *T* = 15 min are analogous, with the difference that for *T* = 20 min there are *K* = 6 groups, and for *T* = 15 min there are *K* = 5 groups. Also, in the two latter cases the maximized MI = log_2_(*K*) is larger than MI calculated under the assumption that all input sequences are equiprobable; moreover, the difference is larger than for *T* = 30 min. These theoretical values are collected in Table 1 and compared with numerical estimates for small noise. One can see that both the maximized MI and the MI calculated by assuming that all sequences are equiprobable agree well with theoretical estimates. Additionally, one may notice that the numerically maximized MI perfectly agrees with the value calculated by assuming that all group representatives have equal probabilities and the other sequences have zero probability.

### Bitrate estimation in the limit of infinitely long sequences

To estimate the upper limit of mutual information for small noise for a given base repeating time *T*, one has to calculate the number of sequences of length *L* that can be accurately transmitted. For *T* = 30 min, the accurately transmitted sequences are those without ‘11’ subsequences. It is straightforward to show that the numbers *nL* of such sequences follow the Fibonacci sequence. Let 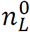 and 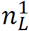 denote the number of such sequences of length *L* ending in respectively ‘0’ and ‘1’. In the first case the sequence can be extended by adding either ‘0’ or ‘1’ at its end; in the second case it can be extended only by adding ‘0’. Thus, we have 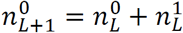, and 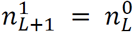 Next, 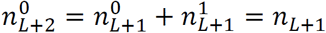 and 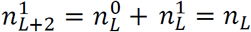, hence *nL***+2** = *nL***+1** + *nL* (i.e. *nL***+1** = *nL* + *nL*-1). Analogously for *T* = 20 min (or any *T* satisfying 2*T* < τ < 3*T*), we may notice that the accurately transmitted sequences are those without both ‘11’ and ‘101’ subsequences. It is also straightforward to show that the number such sequences satisfies a recurrence relation similar to the Fibonacci sequence, i.e., *nL***+1** = *nL* + *nL-2*. Finally, for any *T* satisfying *kT* < τ < (*k* + 1)*T*, the number of accurately transmitted sequences satisfies *nL***+1** = *nL* + *nL*-*k*. In the limit of large *L*, these recurrence relations can be solved: 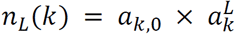 where a_*k*_ satisfies 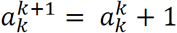, and for *k* ≤ 10, a_*k*,0_ are constants of order of 1: a_1,0_ ≈ 1.17, a_2,0_ ≈ 1.03, a_3,0_ ≈ 0.94, a_10,0_ ≈ 0.74.

Mutual information that can be transmitted in a sequence of length *L* is thus MI(*k*, *L*) = log_2_(*a*_*k*,0_) + *L*×log_2_(*a*_*k*_), thus the bitrate *C*(*k*, *L*) = (*k* + 1)×MI(*k*, *L*)/*L* and in the limit of L → ∞, *C*(*k*, *L*) = (*k* + 1)×log_2_(*a*_*k*_). By straightforward calculation one can find that for *T* = 30 min the bitrate is *C*(2) ≈ 1.39 bit/hour; for *T* = 20 min, C(3) ≈ 1.65; and for *T* = 15 min, C(4) ≈ 1.86 bit/hour. *C*(*k*) is monotonically increasing function of *k*, and *C*(*k*)/log_2_(*k*) → 1 for *k* → ∞.

In Fig 4, for inter-pulse intervals *T* = 60, 30, 20, and 15 min we compare bitrates as a function of sequence length L that were predicted theoretically with those calculated based on numerical simulations. Despite a moderate magnitude of noise (σ = 0.3, µ_0_ = 0.03), for *L* ≤ 8 there is a good agreement between theoretical low-noise prediction and numerical simulations both for not maximized MI (equal input probabilities) and maximized MI. This suggests that the theoretical asymptotic bitrate value, *C*(*k*), may serve as a good approximation even for low and moderate noise, σ ≤ 0.3 and µ_0_ ≤ 0.03.

**Fig 4.**
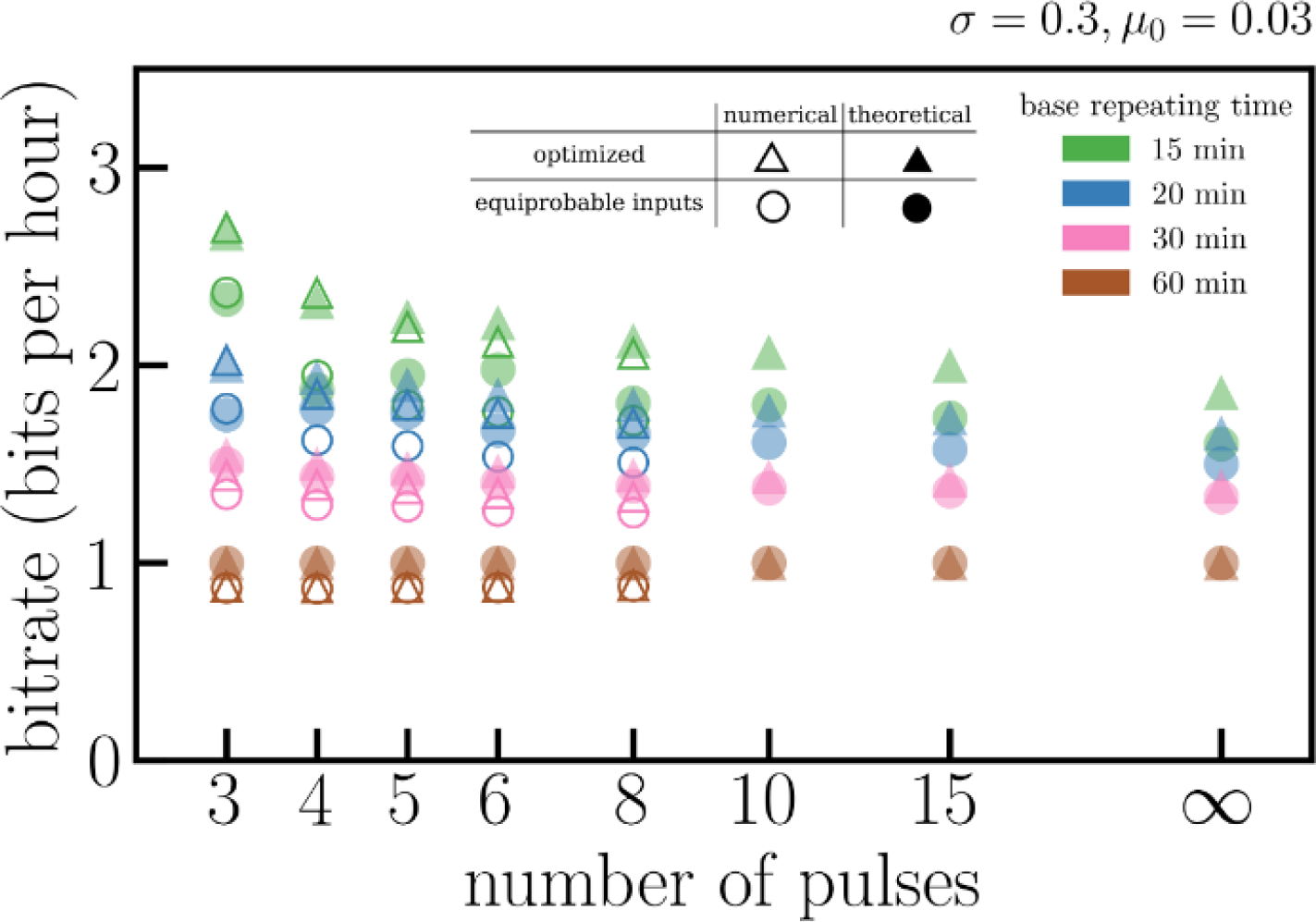
Information transmitted per hour for moderate noise for different carrying frequencies *Φ* = 1/*T*, where *T* = 15, 20, 30, 60 min. Mutual information was calculated as follows: empty triangles – numerically optimized, solid triangles – theoretically maximized, i.e., log_2_(*K*), empty circles – numerically assuming equal probabilities of all input sequences, solid circles – theoretically assuming equal probabilities of all input sequences.

Bitrates increase monotonically with 1/*T*. The maximized bitrate is substantially higher than the bitrate calculated for equal input sequence probabilities. As expected, the difference is most significant for the smallest *T* = 15 min, i.e., when the ratio of sequences that can be properly transmitted to all possible sequences is low. However, even for equal input probabilities the bitrate for *T* = 15 min is much larger than for *T* = 60 min.

### Influence of noise on the bitrate

As one can expect, noise can substantially decrease the amount of transmitted information. As shown in Fig 1D, noise reduces system’s ability to respond to the second EGF pulse, especially when the time span between the pulses is comparable to τ. In Fig 5 we show bitrate with respect to σ, with additive noise kept constant, µ_0_ = 0.03. As expected, bitrates for all considered *T* (60, 30, 20, and 15 min) decrease with σ. Interestingly, the ratio of the optimized bitrate for *T* = 15 min to the bitrate for *T* = 60 min increases with noise, which implies that coding using short base repeating time is less sensitive to noise. This is because in the optimized coding using the short repeating times, the two (non-pseudo) pulses are less frequently separated by 60-min interval than for the base repeating time *T* = 60 min. For *T* = 60 min, subsequences ‘10’ and ‘11’ are equiprobable, and the latter is sensitive to noise (see Fig 1D). Let us notice that even for very high noise parameter σ = 1.0, for *T* = 15 min, *C* > 1 bit/hour (i.e., *C* exceeds the bitrate for *T* = 60 min in the limit of small noise).

**Fig 5.**
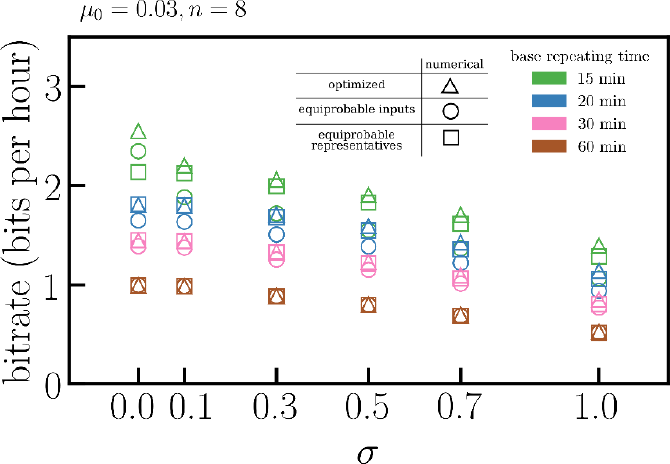
Influence of extrinsic noise on transmitted information. Mutual information was calculated numerically as follows: triangles – maximized, circles – assuming equal probabilities of all group representatives, squares – assuming equal probabilities of all input sequences.

Small noise σ = 0.1 influences bitrate only for *T* = 15 min but more detailed analysis suggests that for *T* = 15 min the high bitrate for σ = 0.0 can be considered an artefact. We found that in this case the system can distinguish between sequences ‘10’ and ‘11’, because the second EGF pulse increases the ERK activity tail. This effect is small and becomes invisible even for small noise (σ ≥ 0.1). The analysis shown in Fig 5 is continued in S1 Fig, where we consider the base repeating times from 30 to 80 min. For σ ≥ 0.3, bitrate increases monotonically with the repeating frequency Φ = 1/*T*; only for small noise, σ ≤ 0.1, is the bitrate for *T* = 60 min somewhat higher than for *T* = 50 min. This effect is clear in the context of Fig 1D, showing that for small noise the pathway transmits pulses separated by 60-min intervals but not by 50-min intervals.

In Fig 6 we repeat the analysis shown in Fig 5, but now analyzing the effect of the additive noise, keeping σ = 0.3. This effect is very weak up to µ_0_ = 0.1 (recall that for µ_0_ = 0.1 the amplitude of the additive noise is equal to 10% of the response amplitude). For µ_0_ = 0.3, the relative reduction of bitrate is significant, and the relative reduction is the strongest for *T* = 60 min. The analysis is continued in S2 Fig for the base repeating times from 30 to 80 min and shows that regardless of the amplitude of additive noise, bitrate increases monotonically with the repeating frequency Φ = 1/*T*.

**Fig 6.**
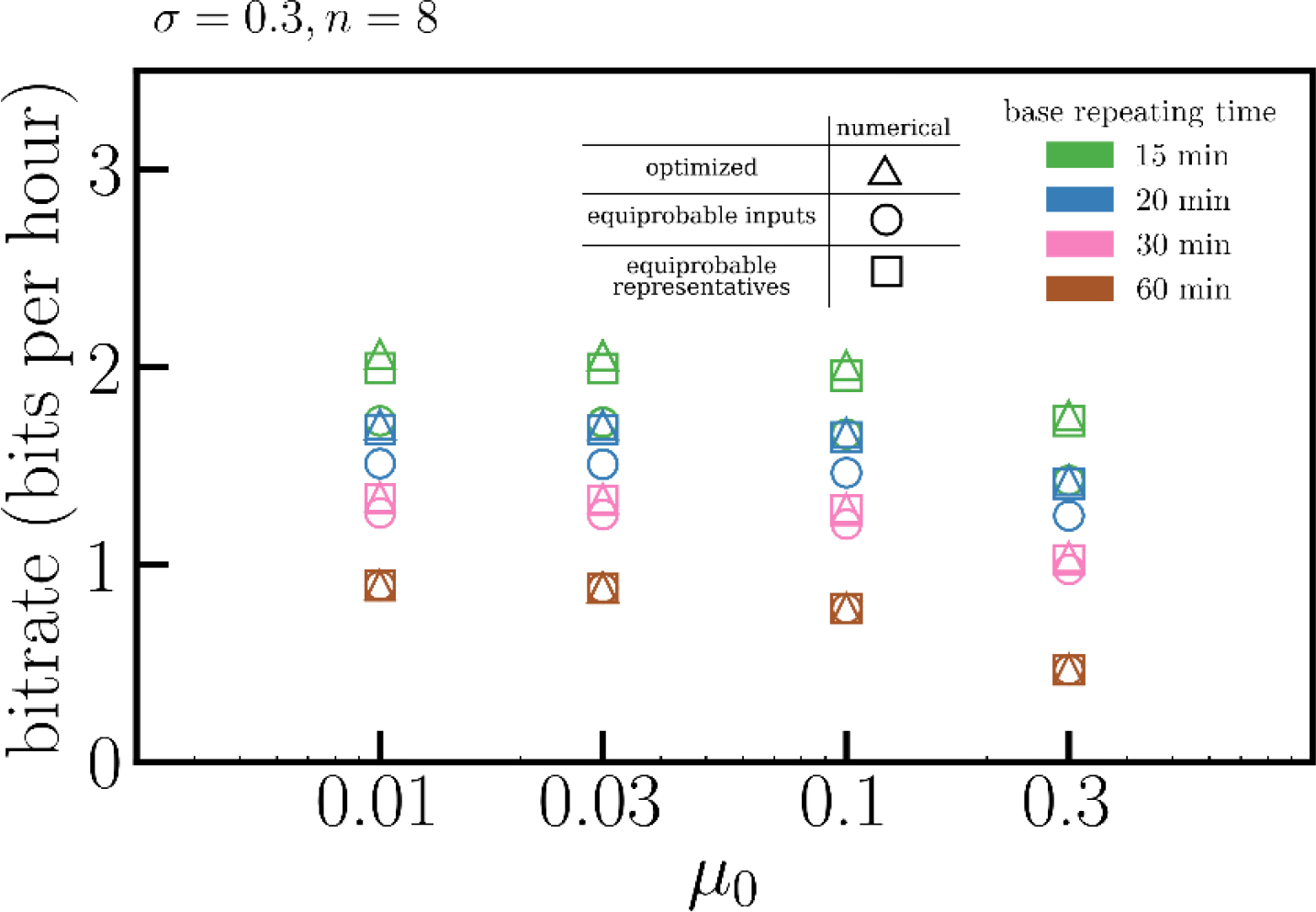
Influence of additive noise on transmitted information. Mutual information was calculated numerically as follows: triangles – maximized, circles – assuming equal probabilities of all group representatives, squares – assuming equal probabilities of all input sequences.

In summary, the analysis shown in Fig 5, Fig 6, S1 Fig and S2 Fig shows that, as expected, noise reduces bitrate. Generically, bitrate increases monotonically with the repeating frequency 1/*T*, for *T* between 15 min and 80 min. Moreover, this increase is steeper for stronger noise. In other words, high-frequency coding is less sensitive to noise than low-frequency coding.

## Discussion

Two important regulatory pathways of NF-κB and ERK were found to transmit merely one bit of information about the level of extracellular stimulation [14,16], which roughly means that these pathways only relay information about the absence or presence of a stimulus (above some threshold). Information about the stimulation dose is increased only slightly when output is measured in more than one time point [16] (see also [18]). This somewhat surprising result may suggest that, in the considered pathways, information is encoded not in the input amplitude but in temporal profile of, respectively, TNF and EGF, that are transcoded to the temporal profiles of nuclear NF-κB or active ERK. The question about optimal information coding can be formulated as the following representation problem: What is the input signal coding allowing to achieve the highest information transmission rate in long time? It has been investigated previously whether the amplitude or the frequency coding can transduce more information from receptors to transcription factors [19]. Here, we propose to code information into sequences of EGF pulses, and based on this we theoretically estimated from below the maximal transmission rate.

The considered model predicts that nearly all EGF inputs are converted to pulses of ERK activity [4]. Relaxation oscillations arise in response to constant EGF stimulation. Oscillation period is a non-monotonic function of the EGF level, attaining minima for the moderate stimulation. Ratio of the ON phase to the OFF phases increases monotonically with the EGF dose; at high dose, oscillations are replaced by constant response [4]. Analogous pulses of EGF are converted to nearly digital pulses of ERK activity. Conversion of various inputs to nearly digital ERK activity pulses my suggest that (at least for some cell lines) pulses are used as symbols in intracellular communication. This motivated us to quantify mutual information between sequences of short EGF pulses and corresponding sequences of ERK activity. Based on this we estimated from below the EGF-to-ERK-channel information capacity, or simply the bitrate at which information can be transmitted.

We showed that high repeating frequency coding allows to substantially exceed bandwidth limit equal 1 bit/τ, where recall τ ≈ 1 hour is the threshold (or relaxation) time. Specifically, we considered sequences of EGF pulses of ‘1’ or ‘0’ amplitude separated by time intervals *T* that are multiples of hour/*n*, and showed that in the case when intervals between pulses of amplitude ‘1’ (true pulses) are not smaller than 1 hour, *C* ≈ 1.39 bit/hour for *n* = 2, and *C* ≈ 1.86 bit/hour for *n* = 4. Because ERK activity pulses are nearly digital, the considered encoding is not very sensitive to noise. Moreover, the high repeating frequency coding (*n* > 1) is less sensitive to noise than coding with *n* = 1, because for the high repeating frequency coding the true pulses (of amplitude ‘1’) are less frequently separated by τ, which is the case when transmission of the second pulse is problematic. The high repeating frequency coding is equivalent to coding by pulse/spike intervals, proposed many years ago for neural information processing [20]. Neuronal spikes are separated by intervals of at least an order of magnitude longer than spike duration [21]. In this case, information is more efficiently encoded by time-intervals between subsequent ‘1’ (that are not shorter than τ and resolved with accuracy τ/*n*) than by sequences of ‘0’and ‘1’ separated by τ.

The bitrate is inversely proportional to threshold time, which for our model is τ = 51.5 min. It should be noted that the value of τ maybe cell line dependent. According to the model, τ decreases with the strength of the negative feedback from ERK to SOS (Fig 7). The abrupt decrease of the τ from 40 min to 25 min is observed when the strength of the feedback is close to 1/4 of its nominal value. At this value, ERKpp does not exhibit oscillations in response to the constant EGF stimulation (Fig 7). Therefore, the model suggests that there is a trade-off between short relaxation time and the ability of the system to translate constant EGF stimulation into series of pulses. In rat PC-12 cells, Ruy *et al*. observed pronounced oscillations in response to 3 min-long pulses of period of 23 min, and also smaller oscillations (that are not arising in our model) in response to the higher frequency pulses [9]. As expected from our analysis, these cells do not exhibit oscillations in response to a constant EGF stimulation. Responses exhibited by PC-12 cell line can be reproduced by the considered model by removing the negative feedback from ERK to SOS and the positive feedback between SOS and RAS. This may suggest that signaling at the level of SOS is different in PC-12 cell line than in human MCF-10A epithelial cells studied by us [4].

**Fig 7.**
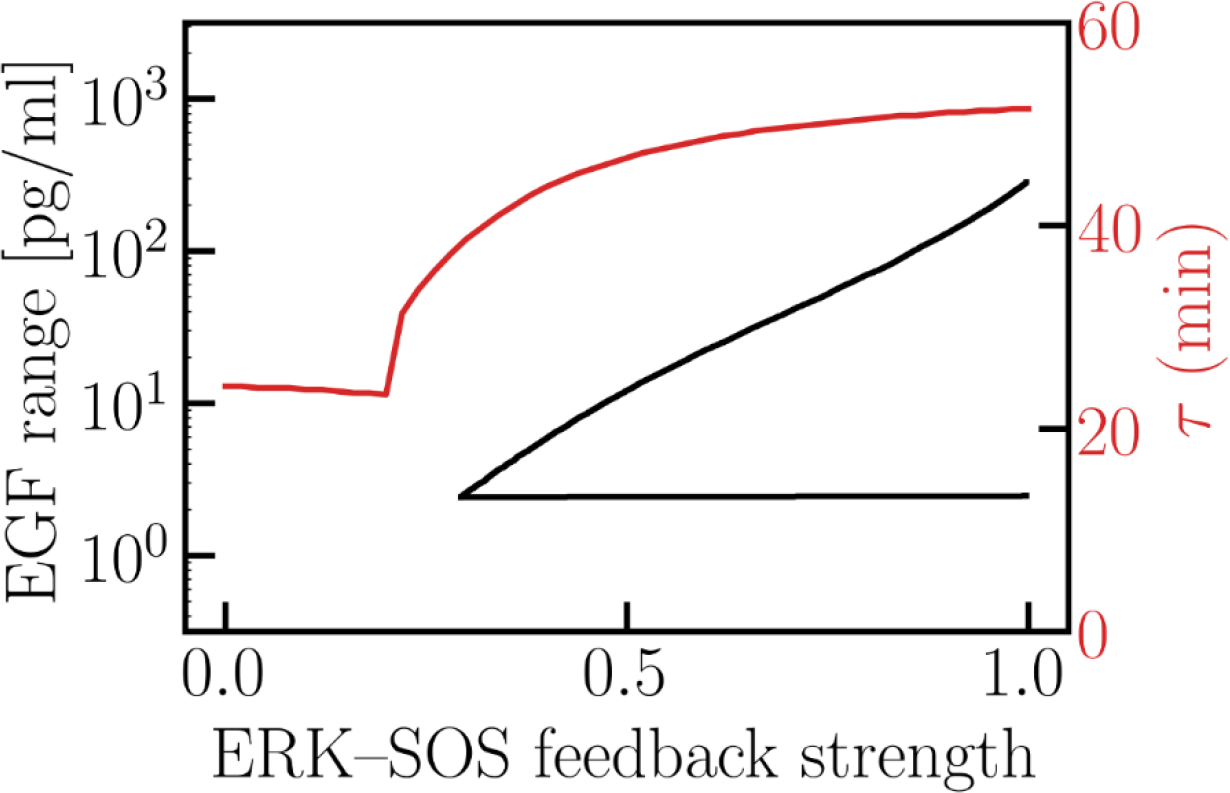
The range of EGF for which *ERK*_*pp*_ exhibits oscillations in response to constant EGF stimulation (black) and influence of the ERK-to-SOS feedback strength on threshold time τ (red).

Characteristic spatiotemporal profiles of cytokines such as TNF and EGF have not been fully characterized. There is however a growing evidence that these profiles are bursty and spatially localized rather than slowly fluctuating and spatially uniform. Capuani *et al*. demonstrated that at physiological EGF concentrations (which does not exceed 1 ng/ml) only a fraction of EGF receptors is activated, which suggests that EGF signaling is short ranged, with EGF being removed from the extracellular space by the receptors [22]. This in turn suggests the existence of spatially localized EGF bursts or waves, that may occur spontaneously [23], and can induce proliferation or motility of selected cells in the tissue. Aoki *et al*. demonstrated that epithelial cells migrate in the opposite direction to the ERK activation waves during wound healing [21]. These activity waves may propagate since ERK activation stimulates a disintegrin and metalloproteases, which cleave membrane-bound ligands of the EGF family. The released EGF-family ligands in turn engage EGFR on the neighboring cells and activate ERK [24]. In the context of NF-κB, recently Bagnall *et al*. showed that also TNF propagation is of short range, and NF-κB signaling is limited to distances of a few cell diameters from the neighboring tissue-resident macrophages [25].

Because the negative feedback loop emanating from ERK targets SOS, a protein at the very beginning of the MAPK cascade, ERK activity oscillations are associated with oscillations of all components between SOS and MEK. Thus same or higher amount of information are reached by pathway components preceding ERK. These proteins control different cellular functions. For example RAF-1, interacts with ROKα implicated in the reorganization of cytoskeletal filaments [26] and with MST2/LATS pathway controlling apoptosis [27].

Regulatory pathways can be perceived as noisy information transmitting channels, but can also be considered as decision making modules that employ feedbacks and other nonlinear regulatory elements to convert input into one of several predefined outputs. MAPK/ERK and NF-κB pathways convert both constant and pulsating cytokine stimulation into pulses of ERK or NF-κB activity. Here, we consider an optimal representation problem and investigate which EGF time-profiles can transmit maximum amount of information to ERK. Motivated by the growing evidence that physiological EGF stimulation is short ranged and bursty, we focus on sequences of short EGF pulses. We found that information can be transmitted with the rate exceeding the classical bandwidth limit of about 1 bit/hour in the case when inter-pulse intervals are used to code information.

## Methods

### Parameters and simulations

To perform our analyses we employed the computational model described in Ref. [4], that has been amended with extrinsic noise and perturbed according to stimulation protocols, characterized by the following parameters:

- pulse duration: 5 min, pulse amplitude: 0 or 100 ng/ml (unless otherwise specified);
- (square of) the second parameter of the lognormal distribution describing additive noise: σ_0_ = 1.0;
- (square of) the second parameter of the lognormal distribution describing cell-specific noise: σ = 0, 0.1, 0.3, 0.5, 0.7, 1.0;
- basic term to determine the first parameter of the lognormal distribution describing cell-specific noise: µ_0_ = {0, 0.01, 0.03, 0.1, 0.3};
- number of pulses: 6 or 8, inter-pulse interval: *T* = 15, 20, 30, 40, 50, 60, 70, 80 min;
- number of simulations for given input sequence of pulses: *M* = 1000.

All parameters of the MAPK model are provided in S1 Table. Abundances of molecules of each pathway component was drawn from a lognormal distribution LogN(µ_i_, σ) with median µ_i_ equal to a default value of this component. Also, all pseudo-first-order parameters, that are proportional to the level of an implicit enzyme, were independently drawn from a lognormal distribution. These randomly set parameters of the MAPK pathway are written in green in S1 Table. For each considered combination of inter-pulse interval *T*, noise strength σ, and specific input sequence of pulses, we generated *M* cell-specific random sets of parameters (as described above) and simulated model dynamics of each cell deterministically using and adaptive ODE solver embedded in BIONETGEN [28]. To simulate the effect of the additive noise of different strengths, we post-processed the obtained *M* trajectories of *ERKpp*(*t*).

### MI calculation and maximization

Estimation of mutual information has been performed as proposed by Kraskov *et al*. [29]. In this approach (see Eq. (8) in Ref. [29]), effectively the output probability distributions, *Y*_i_, and marginal distribution *Y* are estimated based on the distance to the *k*th-nearest neighbor and on the number of points within this radius. For all estimates described in this article, we have set the number of nearestnneighbors *k* = 15, and assumed that distances in input space *X* are much larger than those in *Y* space (which implies that in Eq. (8) from Ref. [28] we set *n*_x_ = *M*). We sampled multidimensional output distributions in *M* = 1000 *d*-dimensional points (the number of dimensions *d* was 2, 3, 4, 5, 6, or 8). We implemented the method of Kraskov *et al*. such that for each input category, *x*, the counts of nearest neighbors from all other categories, *x*’ ≠ *x*, are weighted according to current *p*(*x*’), enabling iterative constrained maximization of estimated mutual information over the complete set of input probabilities. To obtain estimates of the maximum mutual information (channel capacity) and *p*(*x*) associated with inputs, we employed a stochastic gradient optimizer, Adam [30], available in TensorFlow [31]. Our implementation of the channel capacity estimator is provided in the form of a python module called CCE – see S1 Code. The accuracy of our implementation of the Kraskov algorithm is checked within unit tests, where we calculate MI numerically based on 8 three-dimensional Gaussian distributions either (1) centered along an S-shaped curve or (2) centered in vertices of the cube. In both cases the distributions are overlapping so MI is about 1.8 bit. These nearly accurate values can be then compared with the values obtained using Kraskov algorithm basing on *M* points drawn at random from each distribution. Kraskov algorithm overestimates MI by about 2% for *M* = 1000 and 1.5% for *M* = 5000; in the second case, 3.5% for *M* = 1000 and 2.5% for *M* = 5000. Accuracy is nearly the same in the case when all 8 input probabilities are equal (test_use_case_1.py) and in the case when input probabilities are varied to maximize MI (test_use_case_3.py).

## Supplementary information

**S1 Fig. Influence of extrinsic noise on transmitted information.**

**S2 Fig. Influence of additive noise on transmitted information.**

**S1 Table. Parameter values for the MAPK pathway model.**

**S1 Code. Python module Channel Capacity Estimator (CCE).**

